# Predicting forest tree leaf phenology under climate change using satellite monitoring and population-based GWAS

**DOI:** 10.1101/2025.04.30.651385

**Authors:** Markus Pfenninger, Liam Langan, Barbara Feldmeyer, Linda Eberhardt, Friederike Reuss, Janik Hoffmann, Barbara Fussi, Muhidin Seho, Karl-Heinz Mellert, Thomas Hickler

## Abstract

Leaf phenology, a critical determinant of plant fitness and ecosystem function, is undergoing rapid shifts due to climate change, yet its complex genetic and environmental drivers remain incompletely understood. Understanding the genetic basis of phenological adaptation is crucial for forecasting forest responses to a changing climate. Here, we integrate multi-year satellite-derived phenology from 46 *Fagus sylvatica* (European beech) populations across Germany with a population-based genome-wide association study to dissect the environmental and genetic drivers of leaf-out day (LOD) and leaf shedding day (LSD). We show that environmental factors, particularly temperature forcing and water availability, are the primary drivers of LOD variation, while LSD is influenced by a more complex suite of climatic cues. Our genomic analysis identifies candidate genes associated with LOD and LSD, primarily linked to circadian rhythms and dormancy pathways, respectively.

Furthermore, genomic prediction models incorporating these loci accurately reconstruct past phenological dynamics, providing a powerful framework to forecast forest vulnerability and adaptation to future climate change.

## Introduction

Leaf phenology determines when and how long plants photosynthesize, directly influencing whether this activity aligns with favourable environmental conditions. In deciduous trees, leaf phenology links the vegetated land surface with atmospheric energy (e.g. albedo) and gas exchange (e.g. CO_2_ and water vapor), making it a key biological process with significant climate feedbacks^1^. For temperate deciduous trees, phenological timing is also a major fitness trait, balancing the benefits of a longer growing season with the risks of frost or drought^2–4^. Leaf phenology is triggered by environmental cues^5^ but also has a genetic basis^6–10^, recently shown to be complex and polygenic^11^. Like most quantitative traits^12^, it results from both environmental and genetic influences. Climate change is shifting environmental phenology cues in time and space, with major implications for forest ecosystems and climate dynamics^1,13^. With estimates of phenotypic reaction norms and genomic prediction scores accounting for heritable differences, forecasting phenotypes under novel conditions becomes feasible^14^. Accurate predictions therefore depend on disentangling environmental and genetic effects^11,15^. To improve understanding and forecasting of forest responses to climate change, it is crucial to integrate the environmental and genetic dimensions of phenological responses^16^.

Until recently, large-scale, long-term phenotyping of phenological traits was logistically difficult and error-prone^16^. This is now changing with satellite-based remote sensing, which provides spatially and temporally highly resolved phenotypic data suitable for genome-wide association studies (GWAS)^17^. Calibration with field data^18^ has Synthetic Aperture Radar proven effective for capturing forest phenology with high accuracy^19^.

Leaf phenological variation in European beech (*Fagus sylvatica*) has been extensively studied^20–23^; however, most efforts have focused on environmental correlates or candidate gene approaches^24^, leaving the genome-wide basis of phenological adaptation at the landscape scale largely unexplored. Here, we integrate high-resolution satellite-based phenological monitoring with a novel population-based genome-wide association study (popGWAS) ^25^ to investigate the genetic and environmental drivers of spring and autumn leaf phenology. This approach, based on population allele frequencies, perfectly suits remote sensing data, because it does not require individual phenotyping but exploits the mean trait differences among populations^25^. Using a chromosome-scale reference genome, multi-year weather data (2015–2022), and phenological observations across 46 beech stands in Germany, we examine whether this integrated approach can address previous limitations and provide a foundation for large-scale, long-term forecasts of forest responses to climate change.

European beech, a dominant forest species^26^ supporting high biodiversity^27,28^ and recognized by UNESCO for its primeval stands^29^, serves as a model species to demonstrate how coupling large scale phenotyping with population genomics opens new pathways for forecasting ecosystem resilience under future climates.

## Results

### Climatic and Geographic Distribution of Sampling Sites

The 46 sampling sites (Figure 1A) were selected from the core range of *F. sylvatica* (Figure 1B), spanning central climatic conditions of the species’ distribution (Figure 1C). Climatic variation was primarily structured by two gradients: PC1 (44.1% variance) contrasted warm, dry summers with cold, wet winters, while PC2 (27.5%) ranged from wet winters and cool summers to dry winters and hot summers. Sites from the southeastern refugial region ^30,31^overlapped climatically with much of the species’ current range (Supplemental Figure 1A+B).

**Figure 1.**
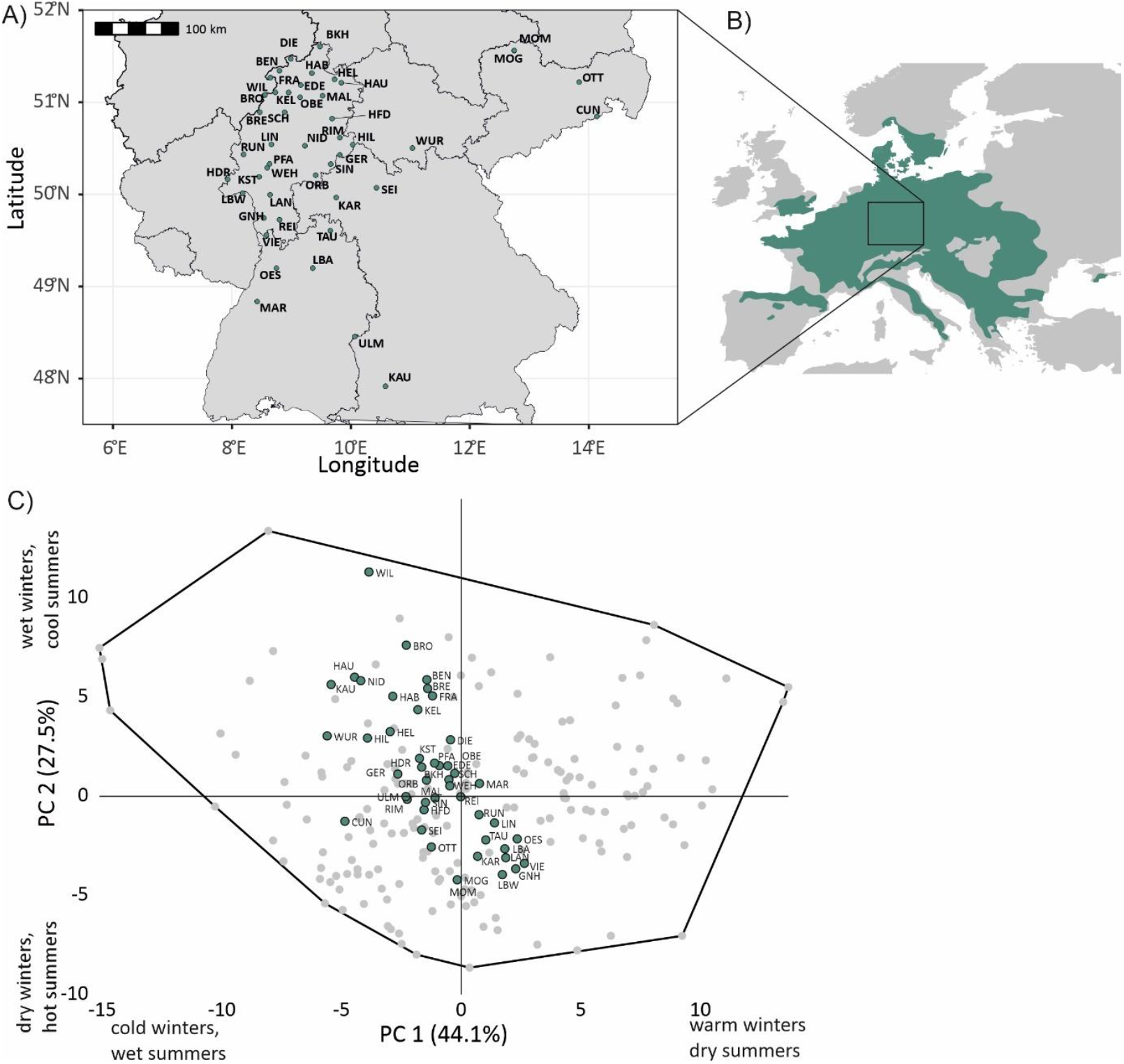
Geographic distribution and long-term climate of *Fagus sylvatica* sampling sites. A) Location of sampling sites. B) Distribution range of *F. sylvatica* in Europe (dark green shading). C) Principal Component Analysis (PCA) of long-term climatic data (1960-1990) of the 46 sampling sites and 178 random points within the species range.

### Remote Sensing of Phenological Dates

Between 2015 – 2022, 62,570 remote sensing observations were collected — averaging 170 per site per year — allowing for near-continuous phenological monitoring. We inferred 339 LOD and 340 leaf-LSD estimates (~ 92 % coverage; Supplemental Figure 2). Five sites (WIL, WUR, GER, ORB, RIM) accounted for most missing data; WIL and ORB were excluded due to insufficient observations (<4).

The mean LOD was 117.13 doy (s.d. 9.57, range 95–146), peaking in late April, and LSD averaged 298.99 doy (s.d. 10.12, range 269–322), peaking in late October. The potential vegetation period spanned an average of 181.72 days (s.d. 15.39, range 137–226 d). Both LOD and LSD varied significantly by year and site (Figure 2A, 2B). LOD was earliest in 2018 (mean 106.79 doy, s.d. 4.74) and latest in 2021 (mean 130.00 doy, s.d. 6.39). VIE had the earliest mean LOD (107.86, s.d. 8.59), and NID the latest (131.14, s.d. 8.9). For LSD, 2017 had the earliest onset (291.71, s.d. 7.39), and 2016 the latest (306.58, s.d. 8.86). BRE showed the earliest average LSD (289.57, s.d. 4.50), and LBA the latest (307.86, s.d. 8.71).

**Figure 2.**
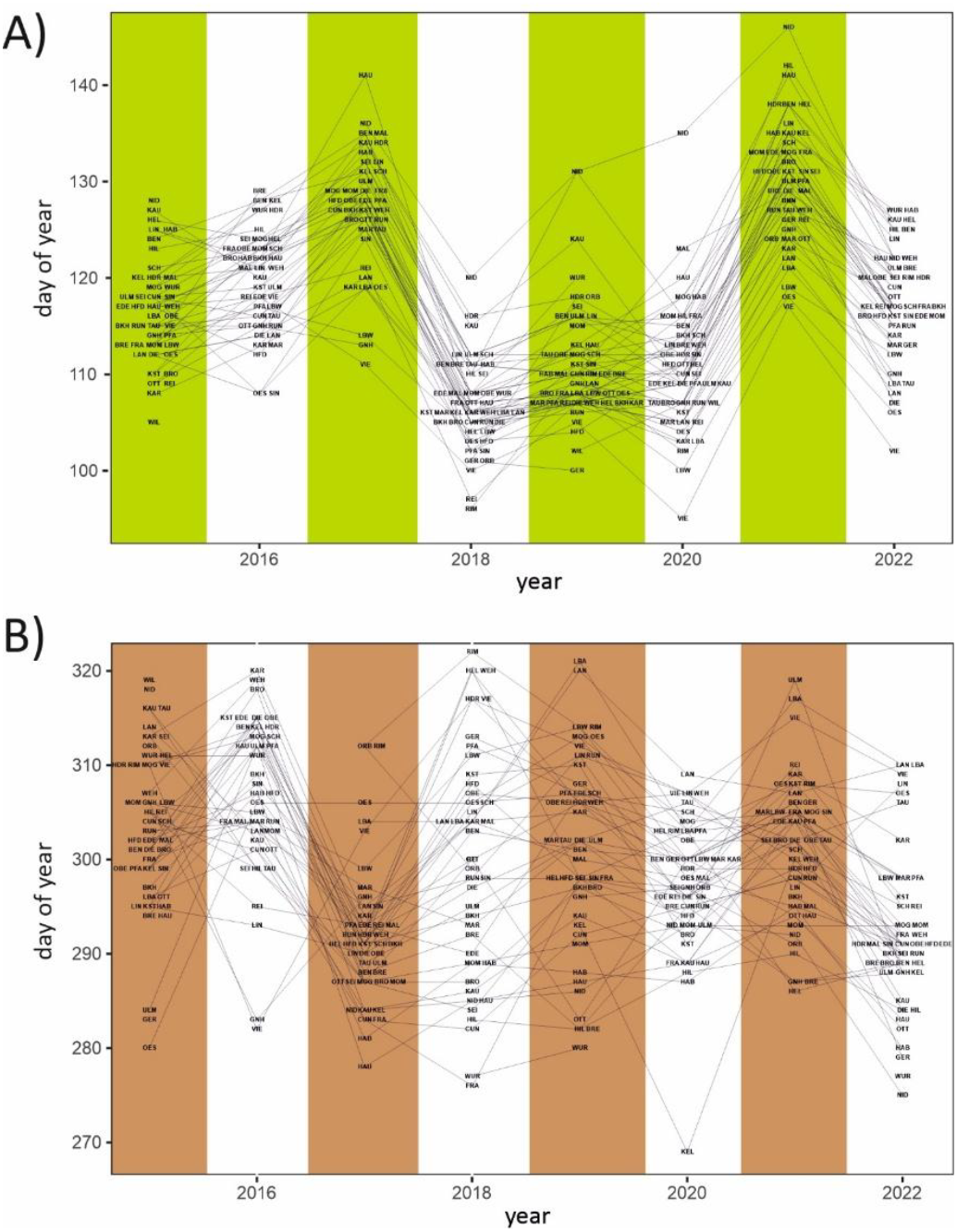
A) Stand-wide mean Leaves-Out-Day (LOD) inferred from Sentinel1 satellite data in the observation period 2015-2022. B) Likewise for Leaves-Shedding-Day (LSD).

### Validation of Remote Sensing Estimates

LOD estimates correlated well with publicly available Germany-wide phenology data (r = 0.79, p = 0.01), lagging by 4.6 days on average (Supplemental Figure 2A). A similar fit was seen between LOD from a phenological garden and a nearby site (LIN, r = 0.77, p = 0.026). For LSD, agreement with national data was moderate (r = 0.57, p = 0.12), with LSD leading by 14.5 days (Supplemental Figure 2C). However, local LSD from the phenological garden did not correlate with the nearby site (r = 0.42, p = 0.29).

### Environmental Drivers of LOD and LSD

To identify environmental drivers of phenological timing, we applied model selection based on previously proposed predictors^8,32–34^ (Supplemental Table 1). For LOD, the best model included frost days, minimum temperatures in January–February, mean temperatures in March–April, and precipitation over those months. This model significantly outperformed alternatives (AIC = 2173; ΔAIC = 26.9; Table 1A), with all predictors having negative coefficients—indicating earlier leaf-out under warmer, wetter conditions. Observed and modelled LOD were strongly correlated (r = 0.757, p < 2.2e-16; Supplemental Figure 4A).

**Table 1.** Results of model selection on phenological variation. A) ANOVA table of best fit model and additional variance explained by systematic site effects on LOD. B) Same for LSD.

| A) Effect | d.f. | SS | p | Coefficient | Interpretation |
| --- | --- | --- | --- | --- | --- |
| Number of days with frost | 1 | 6012 | 7.81e-12 | -0.35 | More frost days trigger earlier leaf flushing |
| Sum of daily minimum temperatures Jan & Feb | 1 | 1150 | $< 2\text{e-}16$ | -0.62 | Warmer winter triggers earlier flushing |
| Sum of daily mean temperature Mar & Apr | 1 | 7400 | $< 2\text{e-}16$ | -1.02 | Warmer spring triggers earlier flushing |
| Mean soil moisture Mar & Apr | 1 | 2385 | 4.39E-07 | -0.10 | More precipitation in spring triggers earlier flushing |
| Residuals | 328 | 13142 |  |  |  |
| Site | 43 | 4227 | 6.92e-13 |  |  |
| Residuals | 287 | 8916 |  |  |  |
| total | 332 | 30090 |  |  |  |

| B) Effect | d.f. | SS | p | Coefficient | Interpretation |
| --- | --- | --- | --- | --- | --- |
| Mean minimum temperature in Oct | 1 | 107 | 0.00387 | -1.15 | Cold October nights induce earlier leaf-fall |
| Number of hot days ($>30^{\circ}\text{C}$) | 1 | 2428 | 7.87e-12 | 0.71 | Hot days in summer prolong vegetation period |
| Mean de Martonne drought index May-Sep | 1 | 56 | 4.45e-06 | -5.30 | Drought in summer induces earlier leaf-fall |
| Mean soil moisture May-Sep | 1 | 3241 | 1.99e-09 | 0.58 | Wetter soil in summer prolongs vegetation period |
| Residuals | 311 | 26374 |  |  |  |
| Site | 45 | 7729 | 2.52e-10 |  |  |
| Residuals | 293 | 19663 |  |  |  |
| Total | 315 | 32205 |  |  |  |

LSD variation was less well captured. The best model (AIC = 2306.9; ΔAIC = 6.5) explained moderate variance (r = 0.432, p = 9.73e-16; Supplemental Figure 4B). Key predictors included mean minimum temperature in October, number of hot days (>30°C), mean de Martonne drought index (May– September), and soil moisture over the same period (Table 1B). Cold autumn nights and dry summers accelerated senescence, while hot extremes and higher soil moisture tended to delay it.

The shortest maximum potential vegetation period averaged over all sites was inferred for the year 2017 (163.71 d, s.d. 11.80), the longest one year later in 2018 (192.59 d, s.d. 14.75; Supplemental Figure 2A).

### Geographic Patterns in Phenology

LOD increased with both latitude and longitude, and was associated with colder, drier winters and cooler, wetter summers (Supplemental Figure 3). LSD showed the opposite trend—decreasing with latitude and longitude, and with colder, wetter winters and cooler, wetter summers. Consequently, the length of the vegetation period varied widely: from just 159.3 days at NID to 210.3 days at RIM.

### Site-Specific Phenological Offsets

On average, sites deviated from predicted LOD values by 2.98 days (s.d. = 2.24). The spread between the earliest (GER, −9.97 d) and latest (NID, +8.23 d) site offset was 18.2 days (Figure 3A). LSD offsets ranged even more: from RIM (+13.54 d) to KEL (–8.60 d), a span of over 22 days. The mean offset for LSD was 3.72 days (s.d. = 3.03; Figure 3B).

LOD offsets weakly correlated with latitude (r = –0.327, p = 0.030), with northern sites flushing earlier than expected (Figure 3C). LSD offsets showed weak positive correlations with long-term winter/early-spring maximum temperatures (r ≈ 0.31–0.32, p = 0.030–0.048), suggesting later leaf-fall under warmer conditions. Notably, LOD and LSD offsets were uncorrelated (r = 0.035, p = 0.82; Figure 3D).

**Figure 3.**
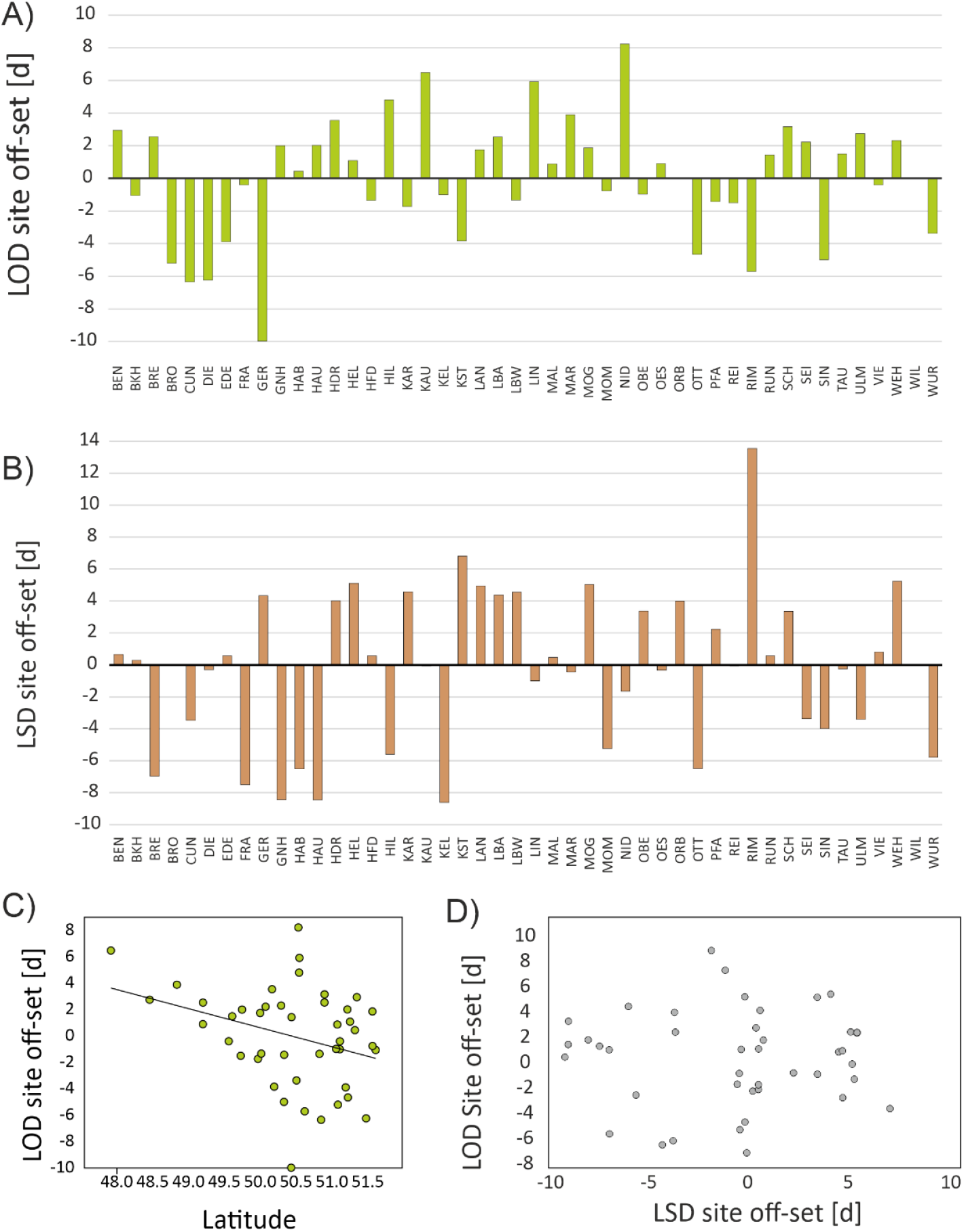
Site specific off-sets in LOD and LSD. A) Site specific off-sets in LOD. B) Site specific off-sets in LSD. C) Plot of the site specific LOD versus LSD effects. D) Plot of LOD site offset against Latitude.

### popGWAS Results

None of the first five PCA axes (explaining together 17.6% of total allele frequency variation as measurement of population structure) correlated with site-specific LOD or LSD offsets (Supplemental Table 2). Combined with low population differentiation (mean FST = 0.062, s.d. = 0.002), this supported the validity of applying popGWAS^25^.

At a 0.9999 threshold (–log10p > 6.0), 22 SNPs were associated with LOD offsets (Figure 4A), spread across all except one chromosome (Chromosome 9, Table 2A). Ten fell within genes, four within 2 kb of genes, and eight in intergenic regions. Among genic SNPs: 3 were intronic, 2 synonymous, and 5 non-synonymous with potentially large protein effects (Supplemental Table 2A). Of the 14 genes associated with these SNPs, 8 were linked to circadian regulation or leaf flushing (Table 2B), while 6 involved retrotransposon elements, or were uncharacterised.

**Table 2.**
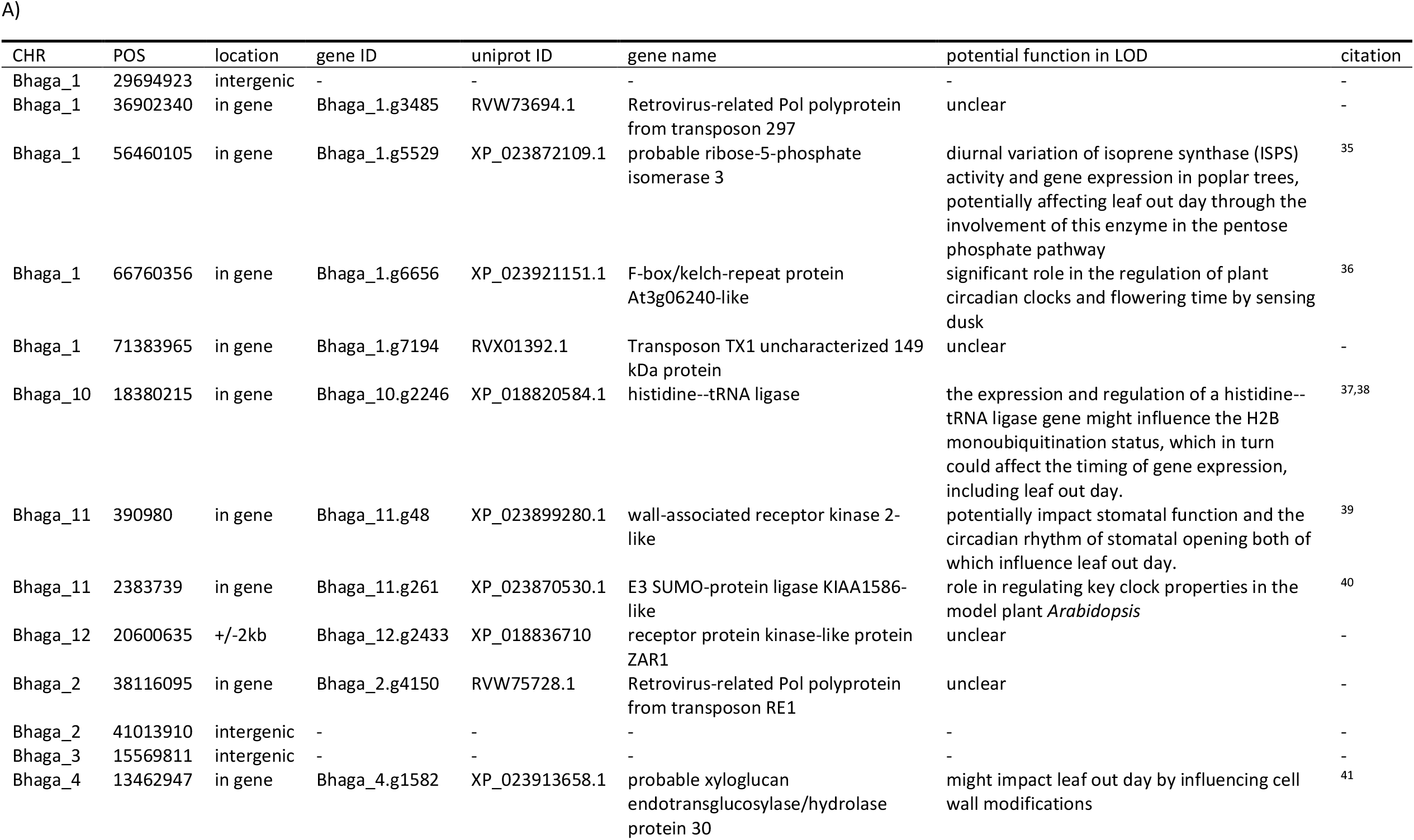
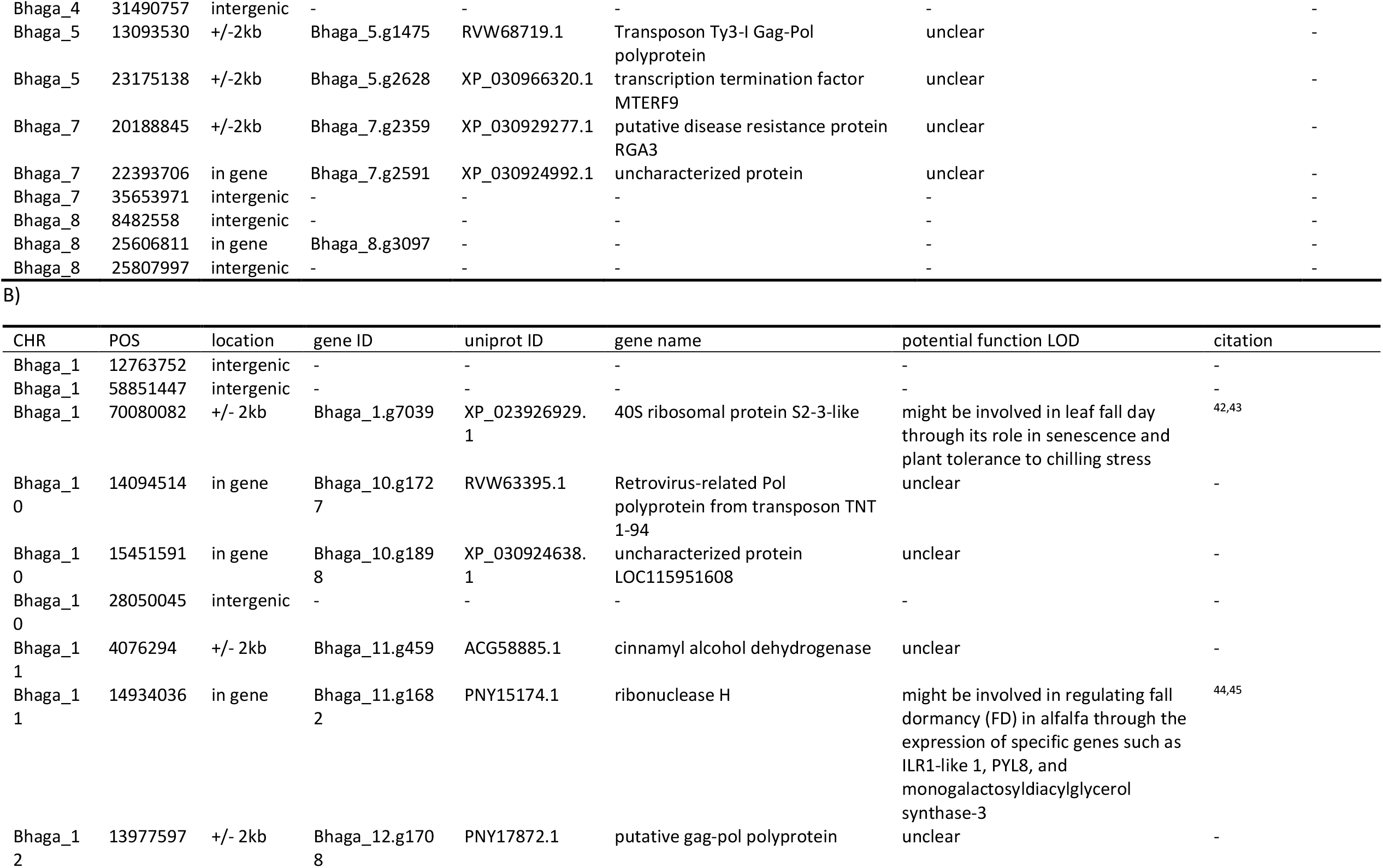
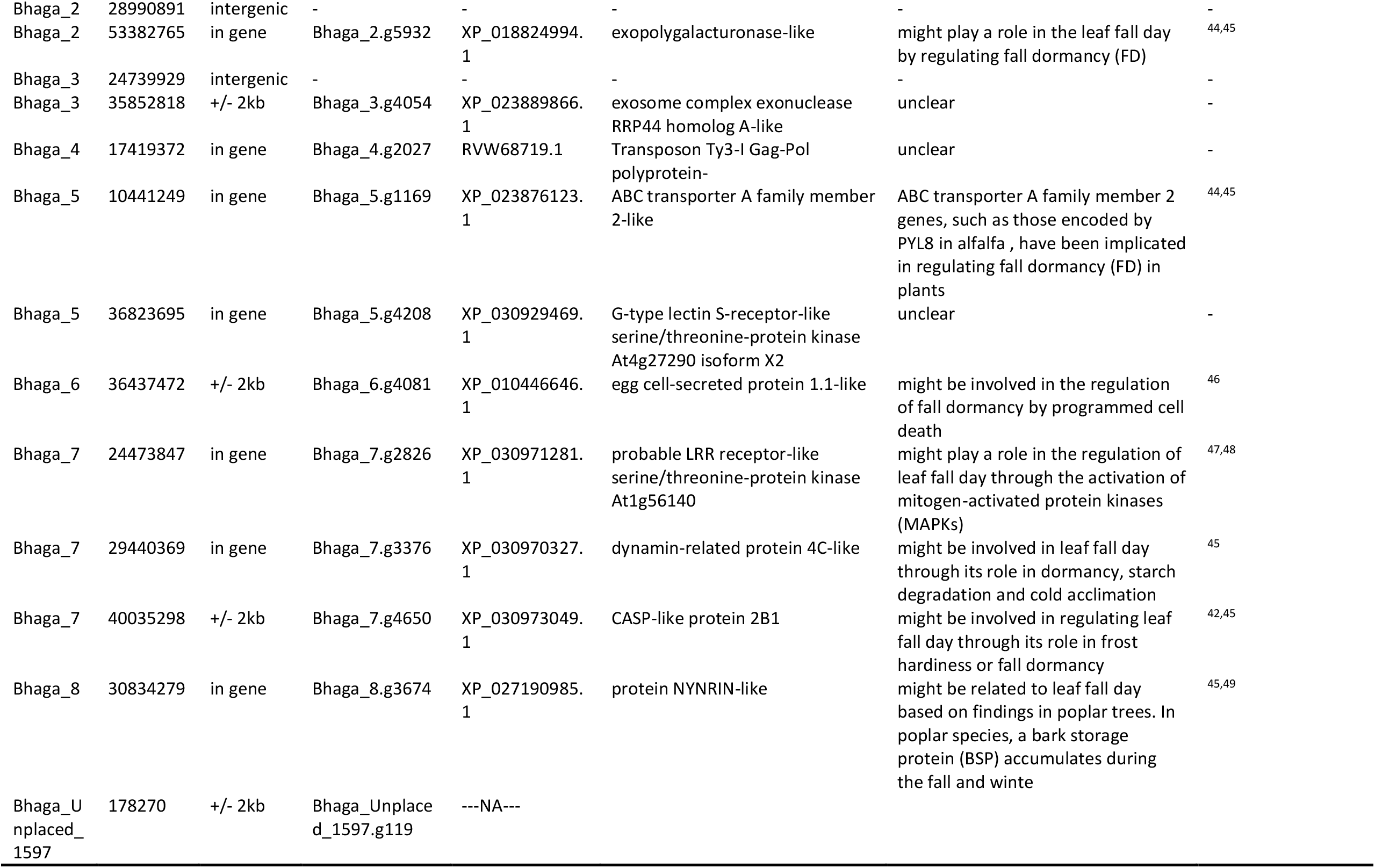
Annotation of outlier SNPs. A) Outlier SNPs for LOD, B) outlier SNPs for LSD. A)

For LSD, 22 candidate SNPs surpassed the 0.9999 threshold (–log10p > 5.2, Figure 4B). Of these, 10 were genic, 7 near genes, and 5 intergenic (Table 4B). Six were intronic, one synonymous, and three non-synonymous (Supplemental Table 2B). Nine associated genes had plausible or known roles in leaf senescence. Transposon-related genes accounted for 4 (LOD) and 3 (LSD) SNPs. For both traits, SNPs on the same chromosome were well-separated (≥1.7 Mb; mean = 14 Mb).

### Genomic Prediction and Functional Validation

LOD phenotypes were best predicted by Minimum Entropy Feature Selection using allele frequencies at 10 selected candidate SNPs (model score > 0.99), achieving 0.97 accuracy in six hold-out populations (Figure 4C). Similarly, 10 SNPs predicted LSD with a score of 0.98 and a validation accuracy of 0.88 (Figure 4D).

Of these ten predictive loci, 7 were polymorphic in 16 independently whole-genome sequenced individuals with available LOD data for the year 2024. Bayesian correlation analysis revealed a 93.9% posterior probability for a positive link between the mean number of LOD-increasing alleles per locus and the observed LOD phenotype, with a moderate median correlation (r = 0.40; 95% HDI: –0.067 to 0.79; Figure 4E).

**Figure 4.**
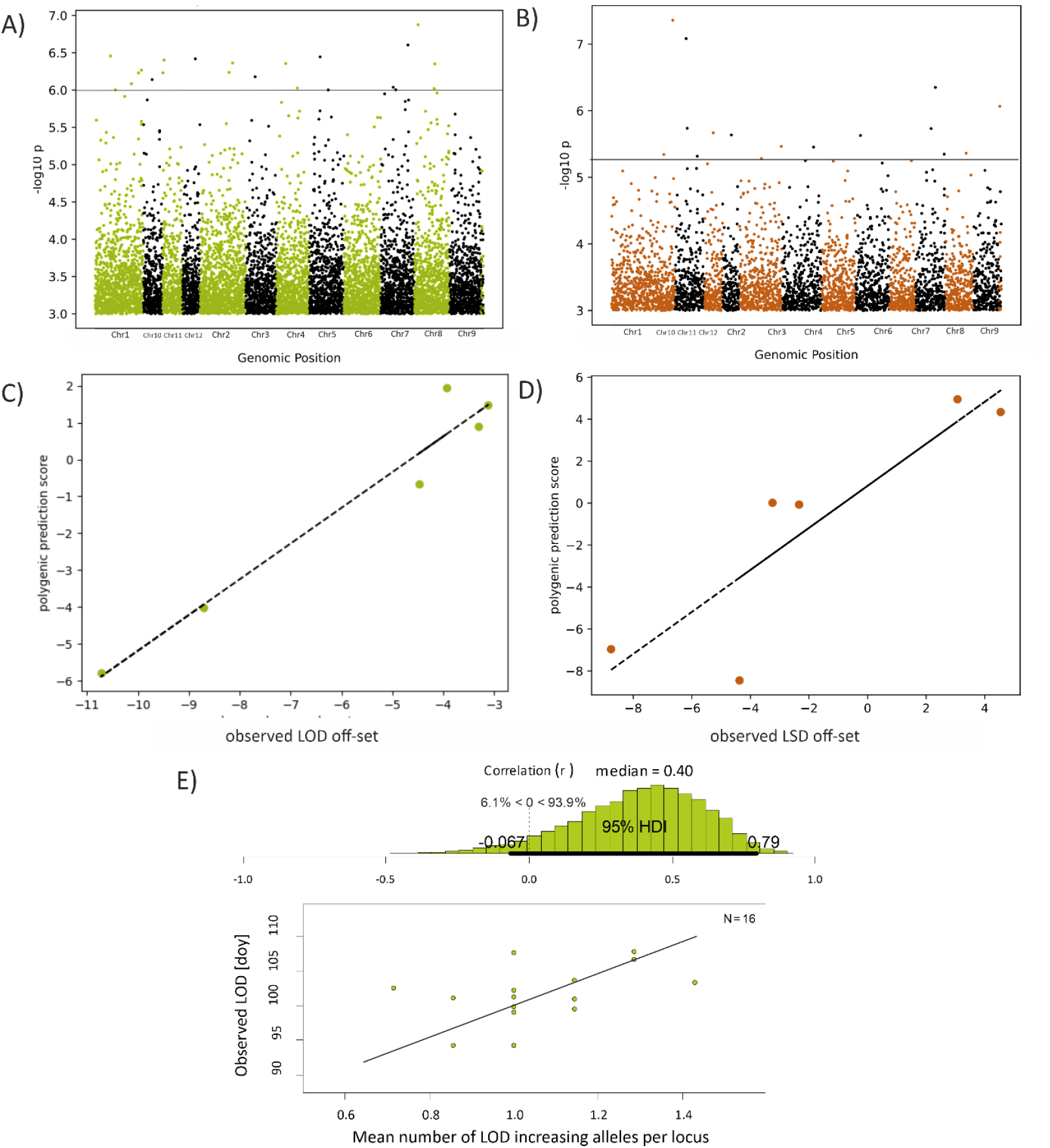
Manhattan plots of popGWAS results and validation of genomic prediction. Shown are the −log10p values above 3.0 along the chromosomes of *F. sylvatica*. A) PLot for LOD. The horizontal line marks the chosen cut-off above which SNPs were considered as candidates. B) The same for LSD. C) Plot of observed LOD off-set scores against the polygenic prediction score derived from hold out populations. D) The same for LSD. E) Functional validation. Bayesian correlation analysis of the mean number of LOD-increasing alleles at predictive loci in 16 individually sequenced trees against their observed LOD date in 2024.

### Modeling Historical Phenological Change

Adding genomic prediction scores to environmental data significantly improved predictive models for both LOD (ΔAIC = 224.7) and LSD (ΔAIC = 138.7), increasing correlations with observed values to r = 0.835 and r = 0.446, respectively. Using these enhanced models to reconstruct phenology between 1971–2022, LOD shifted significantly earlier (slope = –0.16, p = 1.28e–27), advancing by ~1.6 days per decade, or ~8 days total since the 1970s (Figure 5A). While a linear trend described this shift reasonably, a sigmoidal Hill function provided a much better fit (ΔAIC = 416.8, Supplemental Figure 5A). This model identified an abrupt change around 1988, with mean LOD dropping from day 127.4 to 117.6, coinciding with a sharp spring warming (Supplemental Figure 5B). LSD, in contrast, remained stable near the end of October throughout the period (slope = 0.009, p = 0.545; Figure 5B). As a result, the potential vegetation period expanded significantly (slope = –0.22, p = 3.4e–13, Supplemental Figure 6B), increasing from ~167 to ~178 days over five decades.

**Figure 5.**
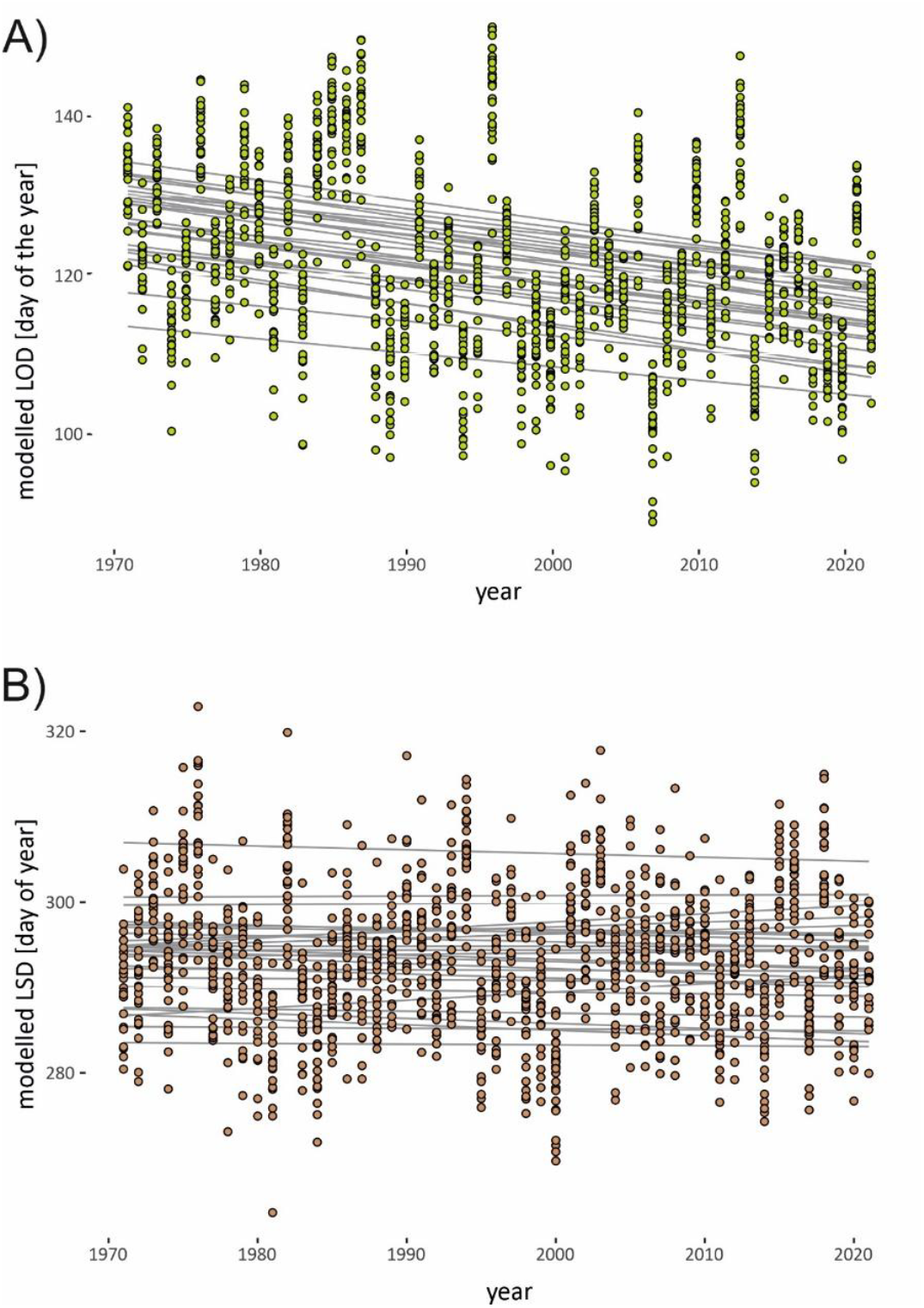
Postdictive modelling of phenological dates for the period 1971-2022. A) LOD modelled for a set of 27 sites. B) LSD modelled for the same sites. The grey lines correspond to the slopes of a linear model fitted for each site.

## Discussion

By integrating satellite-based phenotyping with a novel GWAS framework, we assessed phenological variation across *F. sylvatica* stands with unprecedented spatial and temporal resolution. This allowed us to partition trait variance into environmental and genetic components, identify candidate loci underlying heritable variation, and build accurate genomic prediction models across the species’ core range.

Satellite remote sensing proved highly effective for large-scale phenotyping, enabling consistent leaf status tracking over multiple years—at a resolution that would have required significant logistical and financial investment using field-based methods. Although our primary goal was to capture a reliable, objective signal of canopy-level phenological transitions, prior work^19^ confirms that the strongest change point aligns well with internationally recognized phenological stages. The strong agreement with publicly available phenology data (despite a consistent time lag) supports the reliability of our remote-sensed estimates. Still, limitations exist. Dense undergrowth of beech—particularly where it retains winter foliage—can obscure canopy signals, occasionally preventing accurate LOD or LSD detection. Fortunately, such cases were rare and did not materially impact our analyses.

Both leaf emergence (LOD) and leaf shedding (LSD)—and thus the total growing season—were primarily driven by environmental variation. As expected from previous studies^8,32–34^, early-spring temperature forcing and water availability best explained spatial and temporal variation in LOD, accounting for ~56% of its variance. In contrast, LSD was shaped by competing effects: cold October nights and dry summers accelerated leaf fall, while very hot days and high soil moisture tended to delay it. Although prior work^50^ reported heat as a trigger for early leaf coloring, this may still align with our findings—damaged leaves might senesce earlier but remain attached longer. However, environmental predictors were less effective for LSD. This likely reflects local, stochastic events—like windstorms or heavy rain—that can influence the exact timing of leaf fall but are not captured in large-scale climate data^51^.

LOD and LSD both showed opposing trends with geographical and climatic gradients, suggesting strong environmental control over the vegetation period, which in turn likely shapes beech distribution—particularly at northern and eastern range limits. This complements previous work highlighting drought susceptibility as a key constraint in the east^52,53^. Postdictive modeling supported this interpretation: climate change has already lengthened the growing season at the study sites, mostly through earlier LOD rather than delayed LSD, which is contrary to findings in other studies^54^. The observed advance of ~8 days since the 1970s was not linear. Instead, a sigmoidal model fit the data better, pointing to an abrupt shift in the late 1980s, coinciding with a well-documented surge in spring temperatures^55^.

Both leaf emergence (LOD) and leaf shedding (LSD) showed signs of local adaptation, with likely heritable components varying systematically across space and time. For LOD, site-specific offsets not explained by environmental variables—i.e., the variance where genetic differentiation may reside^56^ —accounted for ~14% of the total variation. These offsets negatively correlated with latitude, suggesting that northern stands initiate leaf-out earlier than expected based on climate alone, likely compensating for photoperiod constraints via local genetic tuning^54^. This supports previous findings of local adaptation in leaf emergence timing^57,58^, and implicates photoperiod perception as a driver^59^.

Site-specific offsets in LSD were even more substantial, explaining ~24% of variance. Stands with generally warmer winters—where frost risk is lower—tended to retain their leaves longer, again hinting at local genetic adaptation. Though this association was relatively weak, it is the first evidence of such a pattern in European beech. Crucially for the GWAS approach, the presumed genetic components of LOD and LSD were entirely uncorrelated, implying distinct genetic architectures for these traits^60^. This independence means natural or artificial selection could potentially act on leaf-out and leaf-drop timing separately^61^. Moreover, predicting the effect of climate change on the length of growing season requires to consider these traits separately.

This study represents the first empirical application of the popGWAS approach^25^. Unlike traditional GWAS, which links individual genotypes to phenotypes, popGWAS uses genome-wide allele frequencies to explain population-level phenotypic variation, offering greater statistical power with the same sequencing effort. Simulations show that popGWAS reliably detects true positive loci for polygenic traits when at least ~36 populations are analysed and a few dozen top outlier candidates are considered. We analysed 46 populations, well above the recommended minimum, and selected 22 top outlier regions per trait as a conservative approach for *F. sylvatica*’s 0.5 Gb genome^25^. A key requirement for successful application is weak population structure unrelated to the traits under study, which was met, as evidenced by the low FST (0.062) and lack of correlation between PCA axes and the genetic components of LOD and LSD.

Many identified outlier loci were located in or near genes known or plausibly related to the traits, making the associations highly unlikely to be due to chance. For LOD, eight genes were linked to the circadian clock or leaf flushing, with similar roles in regulating leaf emergence in tree species like poplar^62,63^. Circadian clock components were shown to be crucial in regulating various growth activities in e.g. poplar trees, including leaf emergence, by modulating gene expression and physiological processes by synchronizing these processes with environmental cues such as light and temperature ^64^. For LSD, nine genes were previously associated with fall dormancy or leaf fall^44,45^. A notable proportion of candidate SNPs were found in transposon genes, where polymorphisms are known to influence gene expression^65^. However, validation and fine-mapping of these loci are necessary to confirm their functional roles^66^. Notably, a positive correlation between the mean number of LOD-increasing alleles and observed LOD in an independent dataset of 16 sequenced beech trees suggests a significant functional role of these SNPs.

We successfully developed accurate statistical models to predict the site-specific genetic component of phenotypes. Prediction accuracy for hold-out populations was high—0.97 for LOD and 0.88 for LSD—despite LSD’s greater environmental sensitivity. This level of success is notable given the modest performance of genomic prediction in other contexts^67,68^. For a correct evaluation, it is essential to distinguish the dual aims of GWAS: identifying causative loci and predicting outcomes. While related, explanation and prediction are distinct. A model can predict well without fully capturing underlying mechanisms, as long as key features correlate strongly with the outcome^69^. Although including more functional loci can enhance predicitve accuracy^25^, it is not necessary to include all or even most trait related loci. Thus, our model should not be interpreted as a comprehensive functional (quantitative) genetic model. Accurate predictions of phenotypic means can emerge when allele frequency differences at just a few loci follow a linear pattern, while other loci behave nearly neutrally. The imperfect fit of outlier loci to linear models reflects the redundancy typical of polygenic traits ^70^. Still, if key loci are shared across populations—via shared ancestry or gene flow—accurate predictions remain feasible. These loci are typically of intermediate frequency, as rare alleles contribute little to variation in population means, regardless of effect size ^25^. Conversely, individual phenotypes may be shaped by rare alleles of large effect^71,72^. Thus, loci important for mean differences among populations may offer limited predictive power at the individual level. Predicting individual phenotypes likely requires a broader set of loci.

Contrary to its application in medicine or selective breeding^68^, however, accurate prediction of population responses is in many instance of the current biodivsity crisis probably more important than the prediction of individual phenotypes^15,73^. Within these limits, our approach yielded highly accurate predictions. While not functionally comprehensive^74^, these prediction scores reliably correlate with trait means and may extend across much of the central *F. sylvatica* range^30,57,75^, given its low genetic differentiation. Such scores could support identifying populations with adaptive potential under future climates.

Our study highlights the power of combining satellite remote sensing with a novel population-based GWAS to assess phenological variation in *F. sylvatica* at an unprecedented scale. Remote sensing enabled consistent, large-scale phenotyping across years, uncovering spatial and temporal patterns in leaf-out and senescence that traditional methods would struggle to capture. By separating genetic from environmental influences, we provide strong evidence for local adaptation and identify candidate genomic regions involved in phenological regulation.

Our predictive modeling further demonstrates the potential of integrating genomic and remote sensing data to forecast population-level phenotypic responses—critical for conservation and forest management under climate change. While functional validation of key loci and refinement of remote sensing techniques remain important next steps, our findings underscore the value of linking high-throughput ecological monitoring with genomic tools to better predict biodiversity responses and inform adaptation strategies for long-lived tree species.

## Material and Methods

### Study populations

We sampled 46 mature beech-dominated stands (>70 years old, >4.5 ha, ≥75% beech basal area) across Germany, emphasizing climatic and ecological diversity with a focus on the state of Hessen. Sites likely included both autochthonous and afforested origins, typical of German beech forests. To minimize sampling of close relatives, 48 canopy trees were collected per site, spaced ≥30 m apart.

### Climatic characterization

Long-term climatic data (1970–2000) were extracted from WorldClim v2.1 at 30″ resolution for all sites and 180 random points across the species range, using DIVA-GIS 7.5^76^ and species distribution shapefiles^77^. We used all BioClim variables and summarized variation via Principal Component Analysis (PCA).

### Remote sensing-based leaf phenology inference

Leaf phenology dates were inferred from Sentinel-1 Synthetic Aperture Radar (SAR) data (VV and VH bands, 10 m resolution), comprising 9236 ortho-corrected scenes (2015–2022) in Interferometric Wide Swath mode^78^, accessed via Google Earth Engine. The VV/VH backscatter ratio was calculated using geemap^79^ (Python 3.11.4), then squared to enhance signal-to-noise ratio. Smoothed time series (LOESS, span = 0.5) were segmented into early and late season (cutoff = day 200). Change points representing maximum signal rate change were estimated using the changepoint package ^80^ (BinSeg, Q=1; R v4.2.2), corresponding to leaf emergence (LOD) and leaf fall (LSD). The period between LOD and LSD was considered the maximum potential vegetation period. Ground-truth leaf phenology data covering Germany from German Weather Service (https://www.dwd.de/DE/leistungen/phaeno_sta/phaenosta.html) and the International Phenological Gardens^81^ (station 189, Linden/Gießen) served as validation, using leaf phenology stage BBCH11^82^ (LOD) and BBCH95^82^ (LSD) as comparison.

### Environmental drivers of leaf phenology

We assessed environmental influences on LOD and LSD using climate data from the German Weather Service (2015 –2022, 1 km^2^ grid, https://cdc.dwd.de/portal/202209231028/searchview). Predictors for LOD included frost/ice days, temperature sums (Jan–Apr), precipitation (Mar–Apr), soil moisture (Mar–Apr), and latitude (photoperiod proxy). LSD predictors included temperature extremes, drought indices (de Martonne), soil moisture (May–Aug), and precipitation (May–Aug). Generalized Linear Models (GLMs) were fit for all parameter combinations; best-fit models were selected using AIC. Site-specific phenological offsets (residuals from the environmental models) were extracted using ANOVA and averaged over years (≥4 years per site required).

### Pooled sequencing and variant calling

From each individual, a 0.5 mm leaf disc (~50 mg) was collected, dried or frozen, and pooled by population. Samples were homogenized (Qiagen TissueLyser), and DNA extracted (Macherey-Nagel NucleoMag Plant kit). Sequencing libraries were prepared and sequenced (150 bp paired-end, ~35x coverage) by Novogene on an Illumina NovaSeq. Reads were trimmed (Trimmomatic v0.39), QC-checked (FastQC v0.11.9), aligned to the beech reference genome (BWA mem v0.7.17), and processed (Samtools v1.10, Picard v2.20.8). PoPoolation2 v2.201 was used to remove indels, calculate allele frequencies, and compute FST values in non-overlapping 1 kb windows (coverage range: 15–50x).

### Population structure and leaf phenology associations

To test for confounding population structure, we performed PCA on allele frequencies, using one SNP per 50 kb with no missing data to reduce linkage. We then assessed correlations between the first five PCA axes and site-specific leaf phenology offsets to verify independence between genetic structure and phenotypic variation.

### SNP–Trait Association (popGWAS)

Using popGWAS^25^, we regressed population mean offset in LOD and LSD against genome-wide allele frequencies. Filtering retained SNPs with ≥15 mean coverage, AF variance ≥0.10, MAF ≥0.10, and ≤35% missing data. For each SNP, we computed linear models and extracted –log10(p-values). Outliers (top 0.01%) were identified and grouped into regions based on 1 kb linkage disequilibrium. The most differentiated SNP per region was retained. A scheme of the pipeline can be found in Supplemental Figure 7.

### Gene Annotation

Outlier SNPs were annotated using BEDtools intersect with the *F. sylvatica* v2 genome GFF^83^. We queried gene functions via UniProt and the ORKG Ask scholarly search and exploration system (ask.orkg.org) with prompts of the following structure “what is known about the role of [gene name] in determining [trait] day?”.

### Genomic Prediction

To predict unmeasured phenotypes, we built a statistical genomic prediction model based on significant SNPs. Missing values were imputed using coverage relaxation, linked SNPs (r > 0.7), or population means. Feature selection was performed via minimum entropy feature selection (MEFS)^84^. The best predictive model was selected using repeated cross-validation over varying feature counts (2 to n–1), implemented in scikit-learn (v1.3.2)^85^. Predictive performance was tested on a hold-out set of six populations using Pearson’s r.

### Candidate SNP Validation

LOD in 2024 was monitored in 14 juvenile beeches at site KST (April 1–29). Individual phenotypes were correlated with the mean number of LOD-increasing alleles via BayesianFirstAid in R. LSD data were unavailable.

### Postdicting Phenological Shifts

We evaluated whether adding genomic prediction scores (SGPS) improved LOD models via AIC. For 27 sites, environmental data since 1971 were used to postdict LOD shifts under observed environmental change using the best GLM + SGPS model.

## Data Availability Statement

The raw data of all samples sequenced can be found at ENA (study numbers: PRJEB64934 and PRJEB60881). Scripts and data at Zenodo.

## Acknowledgments

This study received support from the “Fachzentrum für Klimawandel und Anpassung des Hessisches Landesamtes für Naturschutz, Umwelt und Geologie” in the framework of the FAST-project. This study was partially funded by the Federal Ministry of Food den Agriculture and the Federal Ministry for the Environment, Nature Conservation, Nuclear Safety and Consumer Protection based on a resolution of the German Bundestag (project number 28W-B-4-058-01) and the Bavarian Office for Forest Genetics. We thank Enno Uhl and Matthias Steckel for support in the site and tree selection as well as Yves-Daniel Hofmann, Jorge Cueva-Ortiz and Rebekka Stüwe for data acquisition in the field under sometimes very difficult conditions.

