## Supplementary Info for "Predicting forest tree leaf phenology under climate change using satellite monitoring and population-based GWAS"

**Supplemental Figure 1.** Climate envelopes for species range regions of *Fagus sylvatica*. A) Position of 180 randomly chosen points within the range of *F. sylvatica* and limits of the regions considered. These points were used in conjunction with Worldclim 2.1 data for the period between 1970-2000 with a resolution of 30 sec to construct the climate envelope shown in Figure 1C in the main manuscript. B) Climate envelopes for the seven defined regions. The climate range of the SE region, a Pleistocene refugium of the species, showed the largest range and overlapped to a large extent with all other regions.

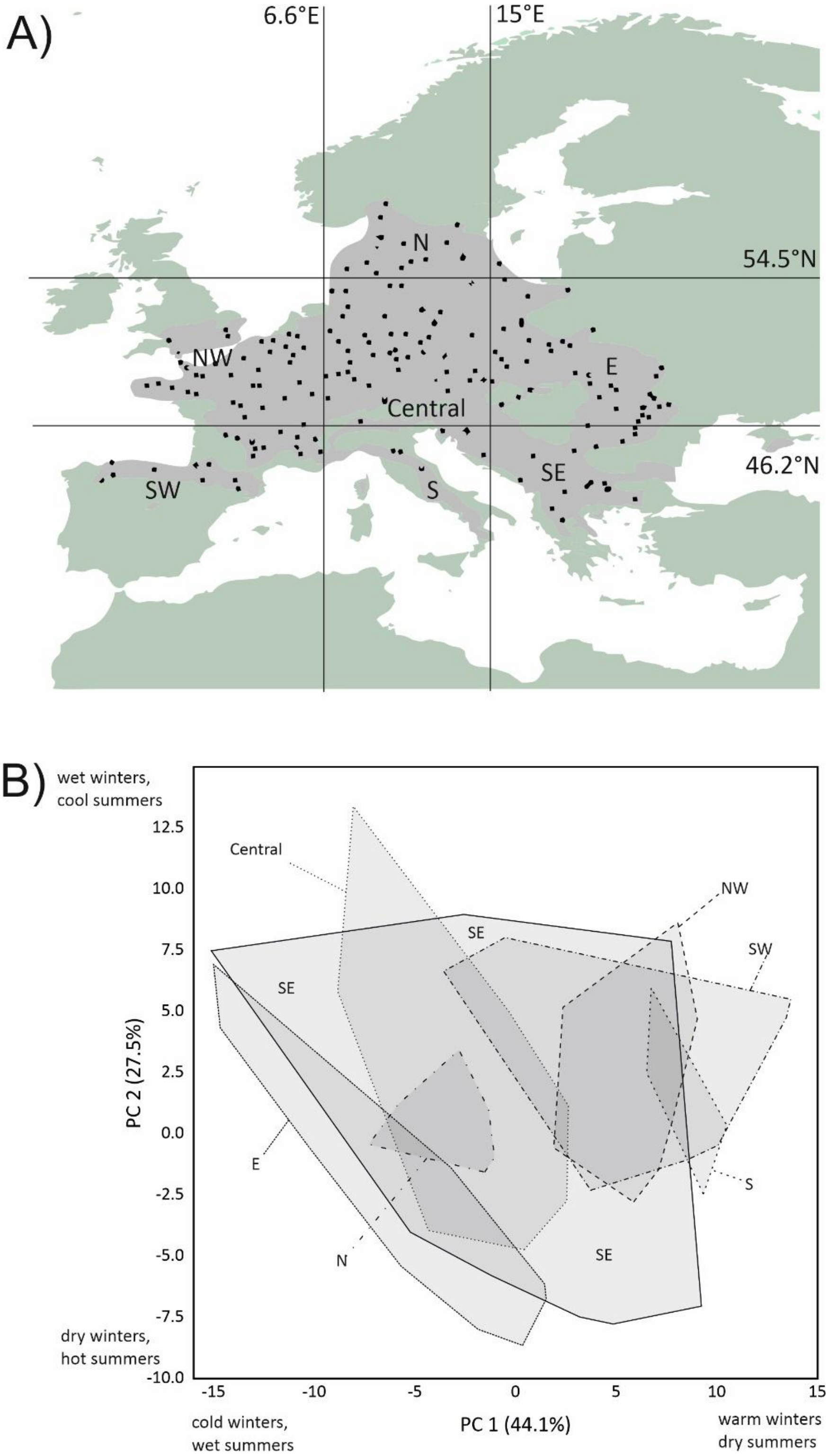

**Supplemental Figure 2.** Scheme of inference of Leaves-Out-Day (LOD) and Leaves-Shedding-Day (LSD). A) Raw stand-wise means of Sentinel 1 VV/VH backscatter ratios for an exemplary stand and year (KST in 2021) plotted against the day of recording. B) Same data, but VV/VH values squared for the first half of the year (light green) and second half of the year (redbrown) with a superimposed LOESS curve (black). The inferred change-point of highest change in the LOESS curve is indicated in orange. C) Data for a site and year (WUR in 2015), where an inference of LOD/LSD was not possible. Please note the continuously high VV/VH values. D) Probable reason for the failure. Natural regrowth of beech tends sometimes to keep the dead leaves during the winter season. Together with high soil humidity, this resulted in a few cases in continuously high VV/VH ratios that did not allow identify LOD/LSD of the respective canopy trees at the site.

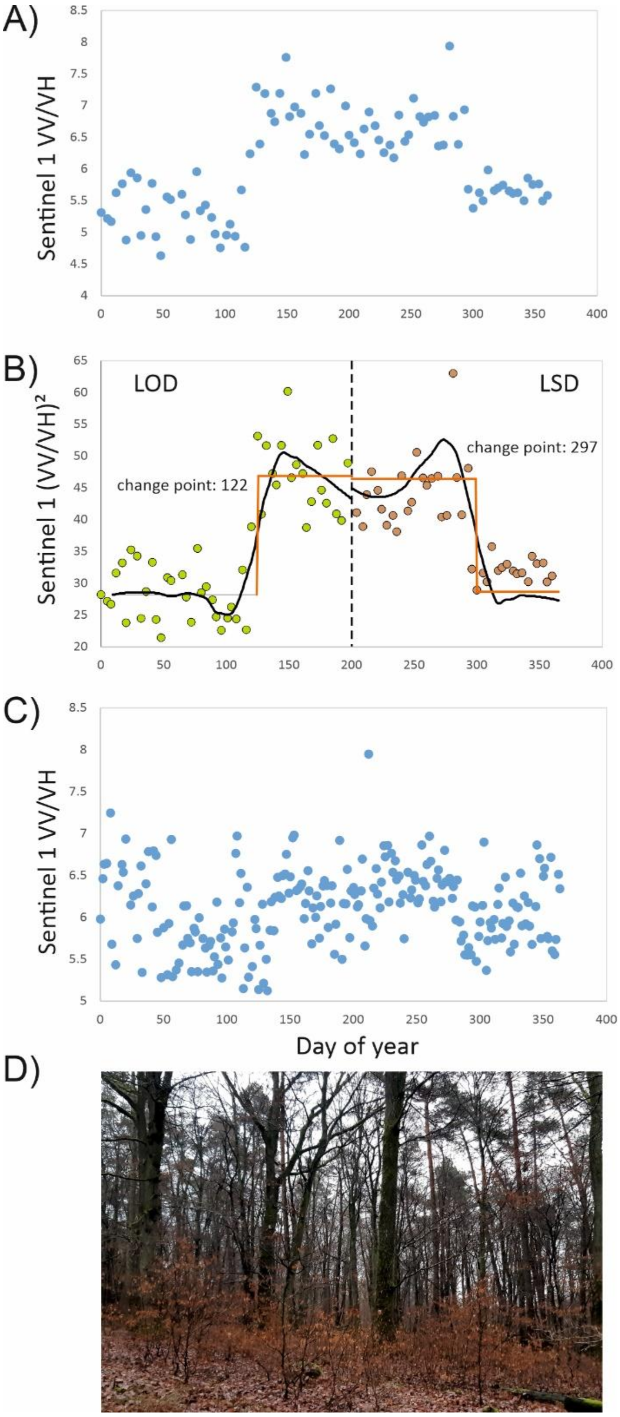

**Supplemental Figure 3**. Validation of RS gained phenological data dynamics with ground observed data. A) Relation between Germany-wide mean BBCH stage 11 day and mean RS observed LOD at all sites (r = 0.85, p = 0.007). B) Correlation between the mean date of the first leaves from four clones planted at the International Phenological Garden Europe in Linden/Hessen and a RS-inferred LOD at a beech forest site (LIN) close to it (r = 0.77, p = 0.026).C) Relation between Germany-wide mean BBCH stage 95 and mean LSD observed (r = 0.17, p = 0.67). D) Correlation between the mean date of the first leaves from four clones planted at the International Phenological Garden Europe in Linden/Hessen and a RS-inferred LSD at a beech forest site (LIN) close to it (r = -0.42, p = 0.29).

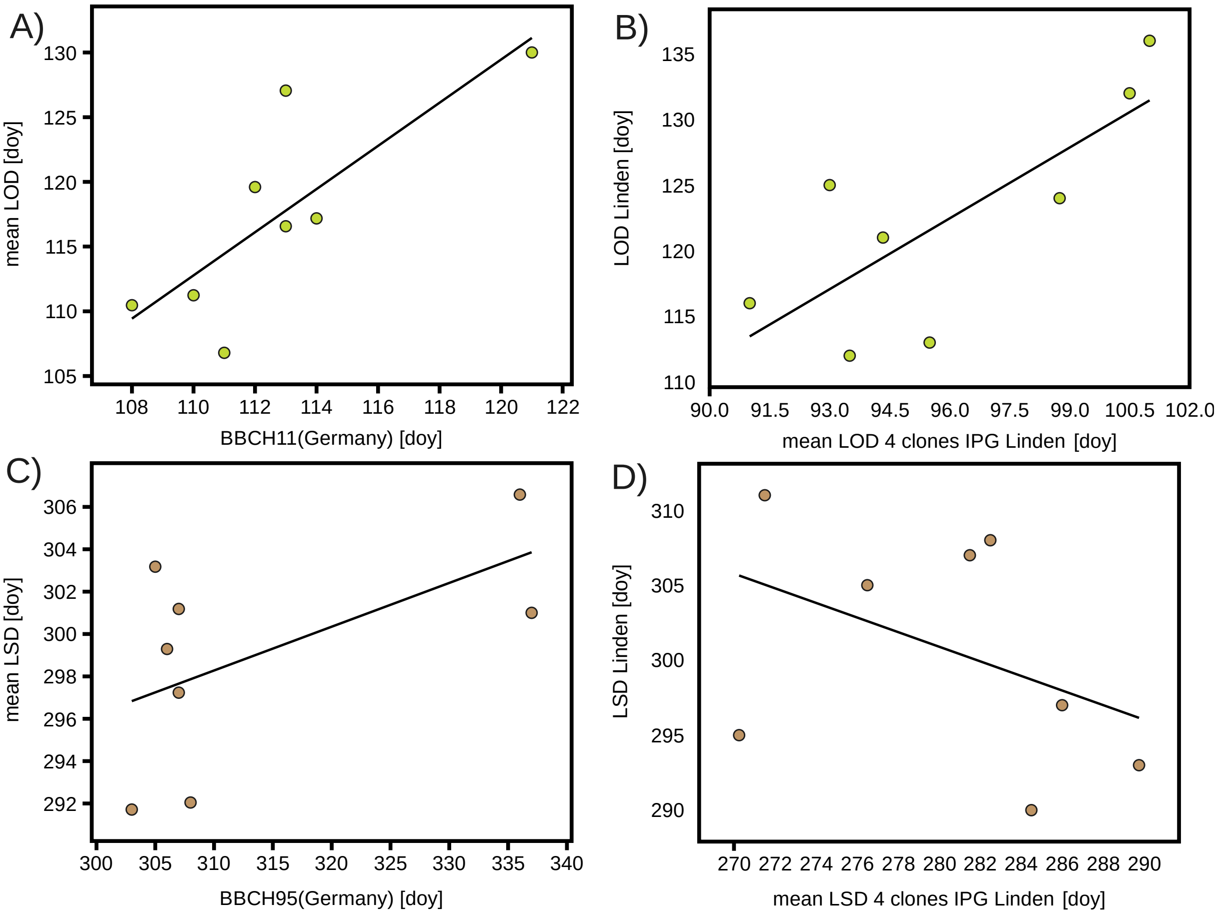

**Supplemental Figure 4.** Correlations of mean phenological dates in the period 2015-2022 with geographical position and long-term climate. 1st row: plot of LOD (light green), LSD (red-brown) and max. potential vegetation period (dark green) against latitude. 2nd row: phenological dates as in previous row against longitude. 3rd row: phenological dates as in first row against climate PC axes 1 (gradient of cold winters with wet summers to warm winters with dry summers). 4th row: phenological dates as in first row against climate PC axes 2 (gradient of dry winters with hot summers to wet winters with cool summers).

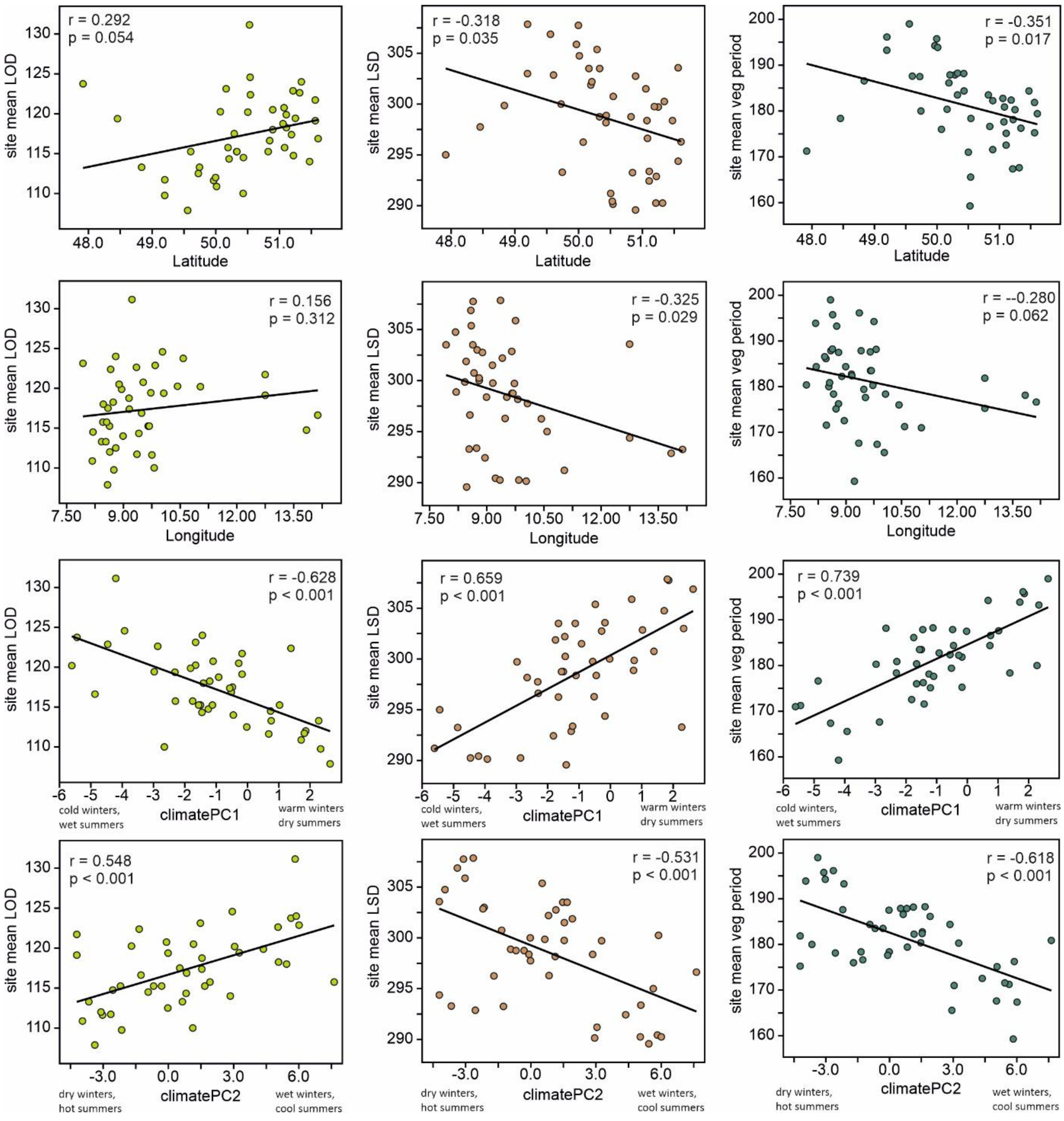

**Supplemental Table 1**. Variables used in model selection. A) Factors potentially influencing leaf-out date (LOD). B) Factors potentially influencing leaf-shedding date (LSD)

A)

| Factor | Variable(s) | Assumed Cause | Literature |
|---|---|---|---|
| Photoperiod | Latitude as proxy | Breaks bud dormancy, avoids late frost | [79–81] |
| Chilling | number of frost days (temperature < 0) | Breaks bud dormancy | [49] |
| | Number of ice days (temperature constantly < 0) | | |
| | Temperature sum mean monthly daily minima in Jan & Feb [°C] | | |
| Temperature forcing | Temperature sum of mean daily temperature in Mar & Apr [°C] | Initiates physiology | [81] |
| Precipitation | Accumulated precipitation in Mar & Apr [mm] | Spring sap rise as precondition | [48,50] |
| | Mean soil moisture in Mar & Apr [%] | | |

B)

| Factor | Variable(s) | Assumed Cause | Literature |
|---|---|---|---|
| Heat | Number of summer days (>25°C) | | [51] |
| | Number of hot days (>30°C) | | |
| Drought | Mean monthly de Martonne drought index Apr-Sep | | [51] |
| | Minimum monthly de Martone drought index Apr-Sep | | |
| | Mean monthly soil moisture May - Aug | | |
| | Minimum soil moisture May - Aug | | |
| Cold | Mean minimum daily temp Oct [°C] | | [51] |
| Precipitation | Accumlated precipitation May-Aug [mm] | | [51] |
| | Mean Soil moisture May-Aug [%] | | |

**Supplemental Table 2.** Effect of candidate SNPs located within the borders of annotated genes A) for LOD and B) for LSD.

A)

| CHR | POS | Ref/Alt | AA exchange/intron | Change in AA characteristics | gene name |
|---|---|---|---|---|---|
| Bhaga_1 | 36902340 | G/A | Alanine > Threonine | Charged to uncharged, impact on secondary structure | Retrovirus-related Pol polyprotein from transposon 297 |
| Bhaga_1 | 56460105 | A/G | intron | - | probable ribose-5-phosphate isomerase 3 |
| Bhaga_1 | 66760356 | G/A | synonymous | - | F-box/kelch-repeat protein At3g06240-like |
| Bhaga_1 | 71383965 | C/T | Proline > Histidine | protein stability and structure | Transposon TX1 uncharacterized 149 kDa protein |
| Bhaga_10 | 18380215 | T/C | intron | - | histidine--tRNA ligase |
| Bhaga_11 | 390980 | G/T | Glycine > Valine | may impact protein conformation and thermal stability | wall-associated receptor kinase 2-like |
| Bhaga_11 | 2383739 | C/T | Aspartic acid > Asparagine | protein post-translational modification, asparagine-linked glycosylation | E3 SUMO-protein ligase KIAA1586-like |
| Bhaga_2 | 38116095 | T/C | Leucine > Proline | protein's structure and thermodynamics, hydrophobicity | Retrovirus-related Pol polyprotein from transposon RE1 |
| Bhaga_4 | 13462947 | C/T | intron | - | probable xyloglucan endotransglucosylase/hydrolase protein 30 |
| Bhaga_7 | 22393706 | T/C | synonymous | - | uncharacterized protein |

B)

| CHR | POS | Ref/Alt | AA exchange/intron | Change in AA characteristics | gene name |
|---|---|---|---|---|---|
| Bhaga_10 | 14094514 | A/G | intron | - | Retrovirus-related Pol polyprotein from transposon TNT 1-94 |
| Bhaga_10 | 15451591 | T/A | intron | - | uncharacterized protein LOC115951608 |
| Bhaga_11 | 14934036 | G/A | Alanine>Valine | similar | ribonuclease H |
| Bhaga_2 | 53382765 | A/C | intron | - | exopolygalacturonase-like |
| Bhaga_4 | 17419372 | G/A | Arginine>Glycine | Neutral to basic, changes in protein stability | Transposon Ty3-I Gag-Pol polyprotein- |
| Bhaga_5 | 10441249 | T/C | Leucine>Serine | structurally dissimilar, secondary structure | ABC transporter A family member 2-like |
| Bhaga_5 | 36823695 | C/T | intron | - | G-type lectin S-receptor-like serine/threonine-protein kinase At4g27290 isoform X2 |
| Bhaga_7 | 24473847 | A/G | intron | - | probable LRR receptor-like serine/threonine-protein kinase At1g56140 |
| Bhaga_7 | 29440369 | T/C | intron | - | dynamin-related protein 4C-like |
| Bhaga_8 | 30834279 | A/G | synonymous | - | protein NYNRIN-like |

**Supplemental Table 2**. Correlation coefficients and significance between site specific off-sets for LOD and LSD, respectively, with the first five PCA axes from a sample of 629 unlinked random SNPs from the genome as measure of population structure.

| | LOD_res | p | LSD_res | p |
|---|---|---|---|---|
| PC 1 | -0.222 | 0.152 | -0.214 | 0.168 |
| PC 2 | -0.275 | 0.074 | -0.225 | 0.147 |
| PC 3 | 0.162 | 0.298 | -0.075 | 0.632 |
| PC 4 | 0.156 | 0.319 | -0.045 | 0.776 |
| PC 5 | -0.071 | 0.653 | 0.254 | 0.100 |

**Supplemental Table 3**. Genomic prediction models for A) LOD and B) LSD. Given are the model coefficients to predict the mean population trait from the allele frequencies at the respective loci.

A)

| LOD | | |
|---|---|---|
| CHR | POS | Coefficient in model |
| Bhaga_1 | 56460105 | -5.078 |
| Bhaga_1 | 71383965 | 8.446 |
| Bhaga_11 | 2383739 | -4.171 |
| Bhaga_2 | 41013910 | -5.573 |
| Bhaga_3 | 15569811 | 7.502 |
| Bhaga_5 | 13093530 | 9.951 |
| Bhaga_5 | 23175138 | -5.643 |
| Bhaga_7 | 20188845 | 8.705 |
| Bhaga_8 | 25606811 | 0.077 |
| Bhaga_8 | 25807997 | 7.634 |

B)

| LSD | | |
|---|---|---|
| CHR | POS | Coefficient in model |
| Bhaga_1 | 70080082 | 11.040 |
| Bhaga_10 | 14094514 | 7.079 |
| Bhaga_10 | 15451591' | 2.219 |
| Bhaga_11 | 14934036 | -19.075 |
| Bhaga_5 | 10441249 | -5.242 |
| Bhaga_7 | 24473847 | -14.454 |
| Bhaga_Unplaced_1597 | 178270 | 2.560 |
| Bhaga_1 | 58851447 | -3.181 |
| Bhaga_6 | 36437472 | 11.939 |
| Bhaga_7 | 40035298 | 8.948 |

**Supplemental Figure 5.** Sudden decrease of mean LOD in the late 1980s. A) Plot of mean modelled LOD for the period 1971-2022.A sigmoid Hill's function (y = 117.58 + (127.37-117.58) / (1+(x/1987.8)^2938.6) provided a decisively better fit to the data (AICc = 4254.9, black line) than a linear model (y = -0.23005x + 580.13, AICc = 4671.7, grey line). B) Annual mean and s.d. of the temperature sum of mean daily temperatures in March and April during the same period. The step-wise behaviour of the modelled LOD is mainly explained by a corresponding sudden increase of the early spring temperatures in the late 1980s.

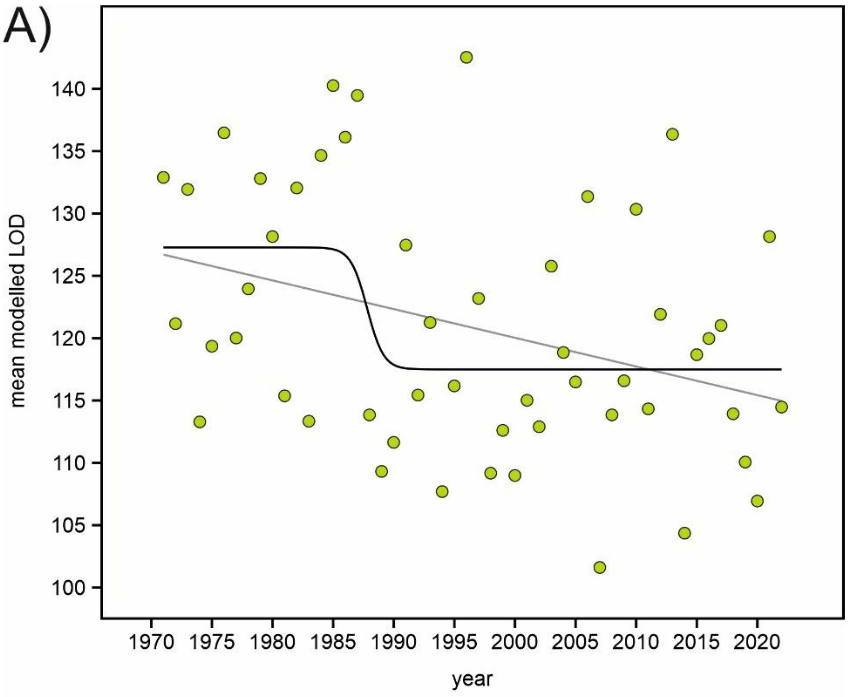


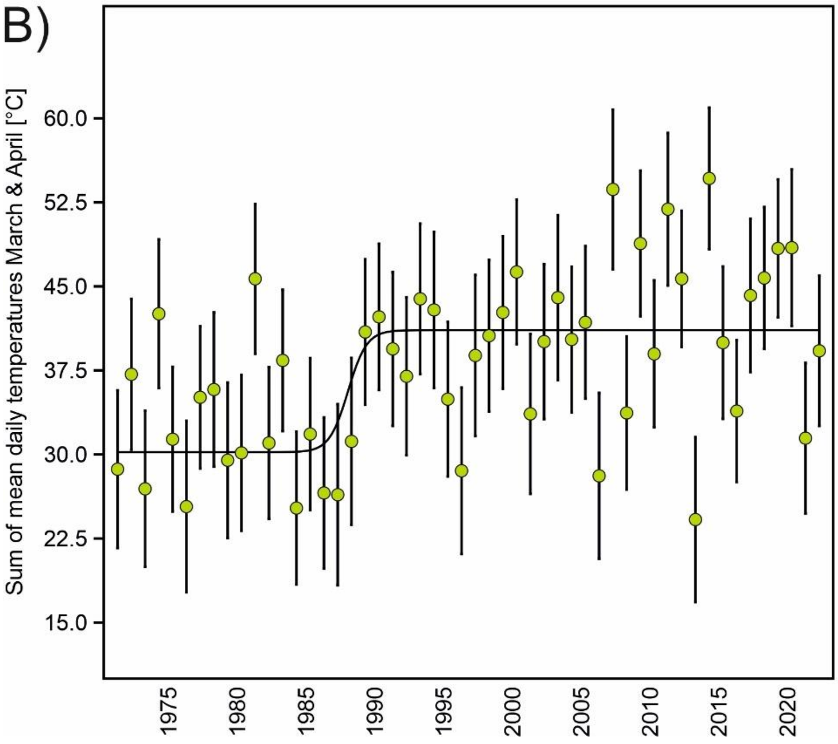

**Supplemental Figure 6.** Results for the inference of the vegetation period length. A) Site-wise length of the maximum potential vegetation period as inferred from observed LOD and LSD dates. B) Postdicted values based on the fitted model and climate data for each site for the period between 1971 and 2022. The light grey lines are the fitted slopes for all sites.

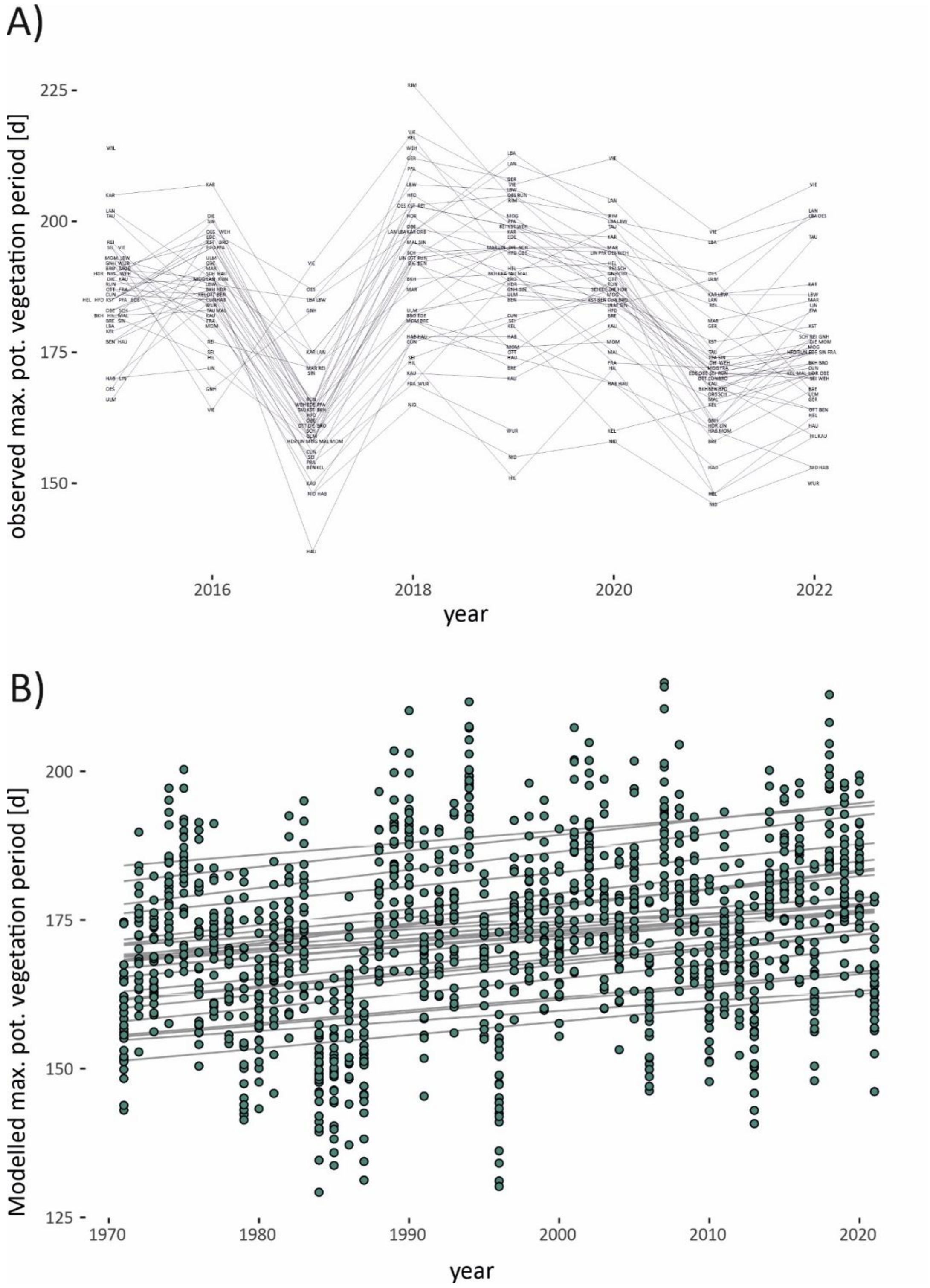

**Supplemental Figure 7**. Schematic overview of the popGWAS pipeline.

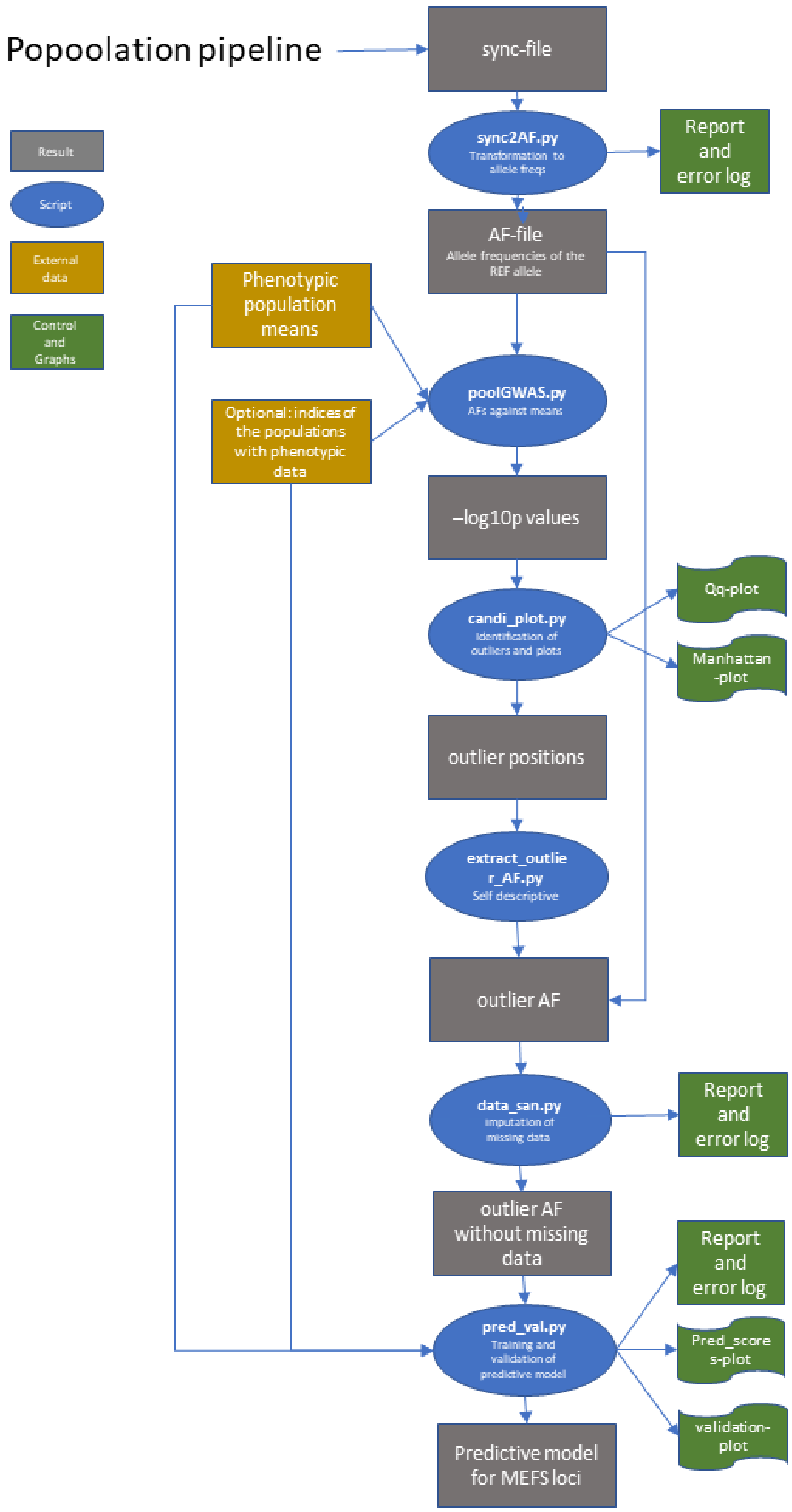
